# Longitudinal study of humoral immunity against SARS-CoV-2 of health professionals in Brazil: the impact of booster dose and reinfection on antibody dynamics

**DOI:** 10.1101/2023.04.03.535504

**Authors:** Ana Paula Moreira Franco-Luiz, Nubia Monteiro Gonçalves Soares Fernandes, Thais Bárbara de Souza Silva, Wilma Patrícia de Oliveira Santos Bernardes, Mateus Rodrigues Westin, Thais Garcia Santos, Gabriel da Rocha Fernandes, Taynãna César Simões, Eduardo Fernandes e Silva, Sandra Grossi Gava, Breno Magalhães Alves, Mariana de Carvalho Melo, Rosiane A. da Silva-Pereira, Pedro Augusto Alves, Cristina Toscano Fonseca

## Abstract

The pandemic caused by SARS-CoV-2 has had a major impact on health systems. Vaccines have been shown to be effective in improving the clinical outcome of COVID-19, but they are not able to fully prevent infection and reinfection, especially that caused by new variants. Here, we tracked for 450 days the humoral immune response and reinfection in 52 healthcare workers from Brazil. Infection and reinfection were confirmed by RT-qPCR, while IgM and IgG antibody levels were monitored by rapid test. Of the 52 participants, 19 (36%) got reinfected during the follow-up period, all presenting mild symptoms. For all participants, IgM levels dropped sharply, with over 47% of them becoming seronegative by the 60th day. For IgG, 90% of the participants became seropositive within the first 30 days of follow-up. IgG antibodies also dropped after this period reaching the lowest level on day 270 (68.5±72.3, p<0.0001). Booster dose and reinfection increased the levels of both antibodies, with the interaction between them resulting in an increase in IgG levels of 130.3 units. Overall, our data indicate that acquired humoral immunity declines over time and suggests that IgM and IgG antibody levels are not associated with the prevention of reinfection.

**Importance:** This prospective observational study monitored the kinetics of humoral response and the occurrence of reinfection in a population of healthcare workers (HCW) who got COVID-19 over a period of 450 days. During the study period, HCW was a prioritized in COVID-19 vaccination campaign, several SARS-CoV-2 variants of concern circulated in the country, and nineteen participants of the study got reinfected. So, we were able to investigate the duration of humoral response against COVID-19, the impact of vaccination boost and reinfection in the production of anti-SARS-CoV-2 antibodies, and the associating of this antibodies with protection against reinfection. These information are important to support health managers in defining COVID19 surveillance and control actions.

## 1. Introduction

Coronavirus disease 2019 (COVID-19), caused by the severe acute respiratory syndrome coronavirus-2 (SARS-CoV-2), emerged in China in 2019, and has affected more than 200 countries. By the end of 2022, there had been 647,972,911 confirmed cases of COVID-19, including 6,642,832 deaths, in the world. In Brazil, the first case was diagnosed in February 2020, and by the end of December 2022, the country had had more than 35 million confirmed cases (1). Healthcare workers (HCW) were a group greatly impacted by the pandemic, leading COVID-19 to be recognized as an occupational disease (2).

Due to the significant impact of the pandemic on health systems, a global effort has sought various alternatives to reduce the harm caused by the disease (3). Thus, vaccines have been developed and approved (4). Due to their higher risk of exposure, HCW were the first group to be vaccinated (5). In January 2021, the vaccination schedule began in Brazil. Since then, more than 80% of the population have had a complete vaccination schedule, and almost 50% have had a booster dose (6). Although these vaccines reduce virus levels in the body, and consequently reduce viral transmission, they do not fully prevent new SARS-CoV-2 infections (7–9). Vaccines have proven to be safe, effective, and timely tools to prevent severe outcomes of COVID-19, including hospitalization and death. However, the efficiency of vaccination can change depending on the type of vaccine and other factors, such as the emergence and/or introduction of new viral variants (9–11).

Since the first recorded cases of COVID-19, new variants of SARS-CoV-2 have been identified. Therefore, in order to establish control and monitoring goals for the new variants, the World Health Organization (WHO) has established three classification categories: variants of concern (VOCs), variants of interest (VOIs), and variants under monitoring (VUMs). The four previously circulating VOCs are Alpha (B.1.1.7), Beta (B.1.351), Gamma (P.1), and Delta (B.1.617.2), while Omicron (B.1.1.529, including BA.1, BA.2, BA.3, BA.4, BA.5 and their descendent lineages) are currently circulating (1). Each SARS-CoV-2 VOC is associated with a new wave of infection, such as the Gamma variant in December 2020 and the Omicron variant in December 2021, which affect human health worldwide (1, 12). Studies involving these new variants have elucidated fundamental aspects of SARS-CoV-2 biology, including viral transmissibility, disease severity, immune system escape, vaccine efficiency, clinical treatment, and management strategies (3, 13, 14).

It is already known that it is possible to become reinfected with SARS-CoV-2 (12, 15). However, since some people have a recurrence of positive test results for SARS-CoV-2 RNA detection during apparently the same infection (16), in order to be considered a reinfection, the CDC has established that the nucleotide sequences of positive samples must be from different lineages or there must be a difference of 90 or more days between positive results (17). Previously published data on the prevalence of SARS-CoV-2 reinfection have highlighted its low level, ranging from 0.10 to 0.65% (18–20). However, the reinfection rate in Brazil is still unknown, especially in risk groups such as health professionals. The emergence of new variants could increase the reinfection rate, as they can escape the immune response triggered by existing vaccines, as with VOCs (e.g., Omicron) (14, 21, 22). Identifying the potential for SARS-CoV-2 reinfection is crucial to understanding the long-term dynamics of the pandemic. Previous studies suggest that the presence of IgG antibodies reduce the risk of reinfection (23). According to a study carried out in England, a primary infection reduces the reinfection rate by 84% over the following seven months (24). Therefore, understanding the dynamic behavior of SARS-CoV-2 infection, and assessing reinfection rates, the impact of genetic variants and vaccines on immune memory kinetics, and their application in the global vaccination campaign are some key points that still need to be elucidated (25).

In this study, we investigated, in HCW in the city of Belo Horizonte in Brazil, the dynamics and longevity of the humoral immune response up to 450 days after the initial onset of COVID-19 disease symptoms and laboratory confirmation of SARS-CoV-2 infection. During the study, the kinetics of the humoral response were monitored both before and after the initial vaccination scheme and subsequent booster dose. Our study demonstrated the occurrence of reinfection in 19 study participants, and showed that (i) the humoral immunity of HCW declined over time, and (ii) the booster dose was essential to increase antibody levels, mainly IgG, but not enough to protect against reinfection with new variants. Robust and constant surveillance is, therefore, essential for responding to future epidemic waves, and provides a basis for recommendations for immunization programs and vaccine updates.

## 2. Results

### 2.1. Description of the study cohort

A total of 163 health professionals aged between 19 to 68 years were included in the initial study cohort between October 2020 to April 2021. Fifty-four (33%) of these individuals subsequently had positive test results by RT-qPCR (Figure 1). Two individuals were excluded from further involvement in the study due to pregnancy or hospitalization, while two other participants withdrew from the study during the follow-up period, but allowed the use of their data and the samples already collected. Therefore, our final study cohort comprised 52 individuals, with 50 of these participants remaining until the end of the 450-day follow-up period. Among the starting 52 participants, four (7.7%) worked at UPA, 16 (30.8%) at HMDCC, and 32 (61.5%) at HC. Overall, the average age of the subjects was 37.38 ± 6.99 years, and 55.8% were female. The final study cohort consisted of 22 (42.3%) physicians, 14 (26.9%) nursing technicians, 10 (19.2%) nurses, and 4 (7,7%) physiotherapists (Table 1).

**Figure 1:**
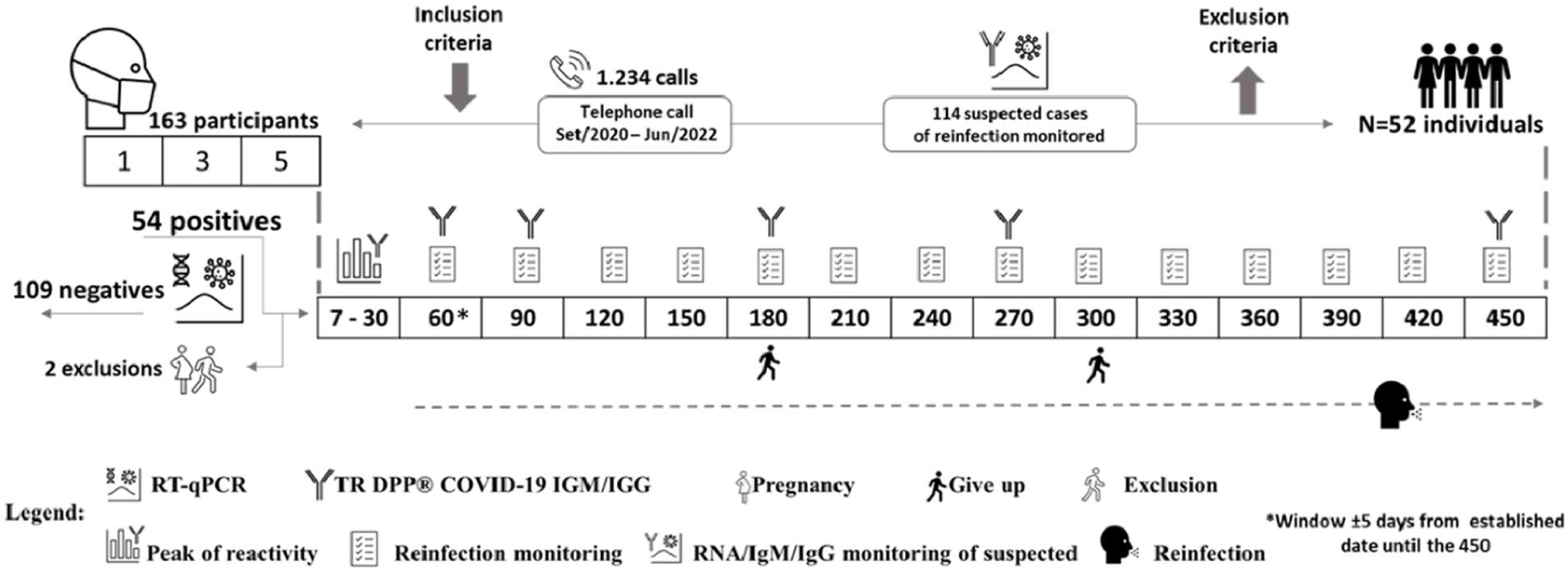
Flowchart illustrating our study workflow. The numbers inside the squares represent the days of visits on which patients were followed-up. The icons shown in the legend represent the study design. After 60 days, questionnaires were answered to assess symptoms consistent with suspected SARS-CoV-2 reinfection, if reported by participants, and clinical samples were collected. Fifty-four participants were confirmed positive for COVID-19, of which one participant became pregnant and another was hospitalized, both were subsequently excluded from the study.

**Table 1.** Demographic characteristics of the final study cohort (*n* = 52).

| Characteristics | Infection | Reinfection | Total | p-value |
| --- | --- | --- | --- | --- |
| Health centers |  |  |  |  |
| HC | 20 (38.5%) | 12 (23.1%) | 32 (61.5%) | 1.000 |
| HMDCC | 10 (19.2%) | 6 (11.5%) | 16 (30.8%) |  |
| UPA | 3 (5.8%) | 1 (1.9%) | 4 (7.7%) |  |
| Total | 33 (63.5%) | 19 (36.5%) | 52 (100%) |  |
| Age |  |  |  |  |
| Range 21-30 | 7 (13.5%) | 5 (9.6%) | 12 (23.1%) | 0.084 |
| Range 31-40 | 7 (13.5%) | 9 (17.3%) | 16(30.8%) |  |
| Range 41-60 | 19 (36.5%) | 5 (9.6%) | 24(46.2%) |  |
| Total | 33 (63.5%) | 19 (36.5%) | 52(100%) |  |
| Sex |  |  |  |  |
| Female | 14 (26.9%) | 15 (28.8%) | 29 (55.8%) | 0.011 |
| Male | 19 (36.5%) | 4 (7.7%) | 23 (44.2%) |  |
| Total | 33 (63.5%) | 19 (36.5%) | 52 (100%) |  |
| Professional Category |  |  |  |  |
| Physicians | 15 (28.8%) | 7 (13.5%) | 22 (42.3%) | 0.691 |
| Nurse Technician | 7 (13.5%) | 7 (13.5%) | 14 (26.9%) |  |
| Nurse | 6 (11.5%) | 4 (7.7%) | 10 (19.2%) |  |
| Physiotherapist | 3 (5.8%) | 1 (1.9%) | 4(7.7%) |  |
| <sup>1</sup>Others | 2 (3.8%) | 0 (0.0%) | 2 (3.8%) |  |
| Total | 33(63.5%) | 19 (36.5%) | 52 (100%) |  |
| Complete vaccination scheme (i.e., initial two doses only) |  |  |  |  |
| Yes | 5 (9.6%) | 19 (36.5%) | 11 (21.2%) | <0.001 |
| No | 28 (53.8%) | 0 (0.0%) | 41 (78.8%) |  |
| Total | 33(63.5%) | 19 (36.5%) | 52 (100%) |  |

<sup>1</sup>Professionals included in the study: social workers, nutritionists, and psychologists.

In January 2021, the vaccination campaign started in Brazil. Thus, 30 (58%) participants were infected with SARS-CoV-2 before starting the vaccination schedule, 11 (21%) during the schedule interval, and 11 (21%) after the complete vaccination, that is 15 days after the second dose of the initial vaccination scheme. During the 450 days of follow-up, 46 (88%) participants had suspected reinfections; of these, 19 (37%) cases were confirmed by RT-qPCR (Figure 2). The proportion of women who became reinfected was higher than that of men (p=0.011). All participants who were reinfected were vaccinated (p<0.001), and 11 had the booster dose.

**Figure 2:**
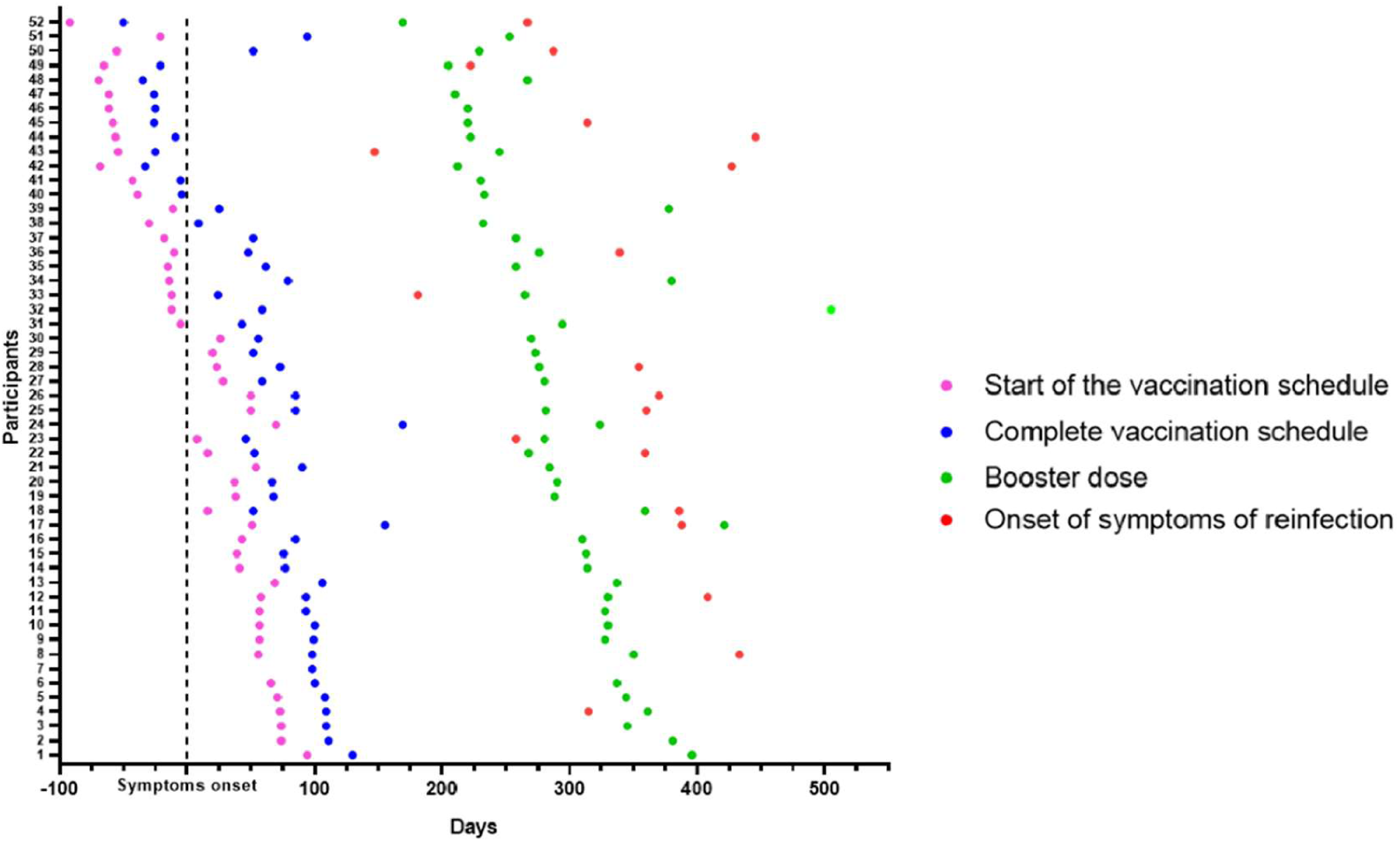
Overview of participant cohorts. Longitudinal follow-up of 52 participants who had COVID-19 confirmed by RT-qPCR. The day of symptom onset was called 0 (Black stroke Symptom onset). The longitudinal timeline shows the days of the start of the vaccination schedule (pink dots), of the complete vaccination schedule, considering 15 days after the second dose (blue dots), and of the booster dose (green dots). Nineteen participants had confirmed SARS-CoV-2 reinfection (red dots).

### 2.2. IgM and IgG antibodies against SARS-CoV-2

The antibody peak observed between the seventh and thirtieth day (D7-D30) of each participant was chosen to be included in the analysis and represented a reaction average of 72.6 (±68.0) for IgM (Figure 3A) and 193.7(±121.2) for IgG (Figure 3B). Nine (17.3%) of the 52 participants did not IgG seroconvert within 30 days, and two (3.8%) individuals did not seroconvert at any time during the study to either of the two monitored immunoglobulins. In addition to these two latter individuals, 12 (23%) did not seroconvert to IgM throughout the study. The reactivity rate for IgM varied from 65%, 36%, 24%, 10%, 15%, and 24%, (Figure 3A) whereas for IgG it was 83%, 85%, 90%, 76% 58%, and 72% (Figure 3B) on days D7-30, D60, D90, D180, D270, and D450, respectively. Immunoglobulin levels decreased over time, reaching their lowest level on day 270 after infection when the mean level for IgM was 13.8 (±17, p <0.0001) (Figure 3A) and for IgG was 68.5 (±72.3, p<0.0001) significant reduction in comparison to the values observed at D7-D30. Figures 3C and 3D show the individual profile for anti-SARS-CoV-2 IgM and IgG in the study participants, respectively. There was no difference between biological sex in the dynamics of antibodies (Figure 4: A and B). However, significant differences in the dynamics of antibody levels were observed, with older individuals presenting the highest levels of IgM and IgG (Figure 4:C and D) and a slower decrease in anti-SARS-CoV-2 IgG levels (Figure 4:D) compared to IgM (Figure 4:C). No significant differences in the Ct value obtained in the RT-qPCR at baseline were observed between the different genders and the age groups (Figure 4: E, F). Although the mean levels of anti-SARS-CoV-2 IgG antibody did not differ significantly between days 270 and 450, representing the period when the majority of reinfection and vaccination booster doses occurred, a tendency for IgG levels to increase was observed (Figure 3: A-F).

**Figure 3.**
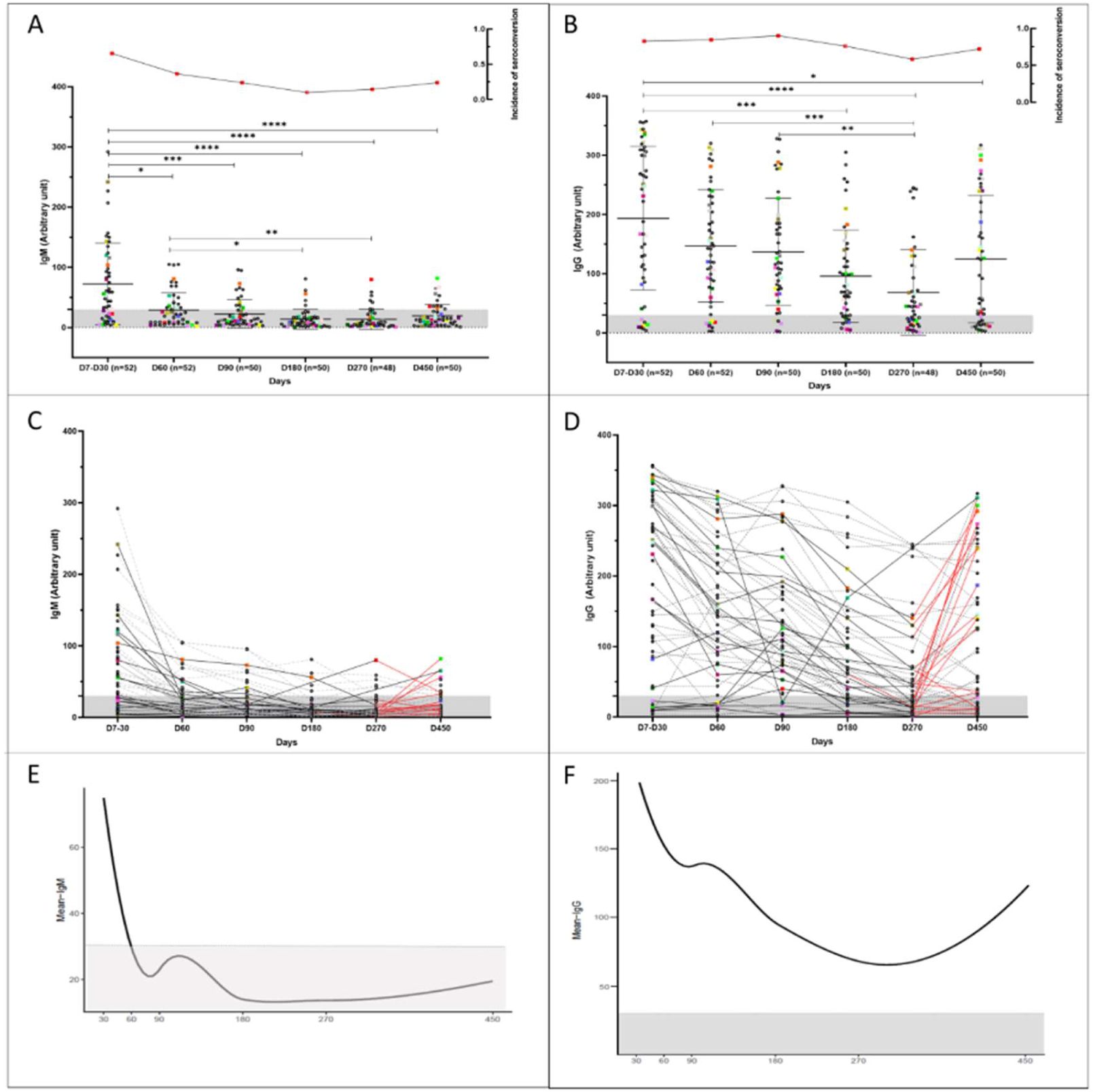
Longitudinal humoral immune response in participants infected with COVID-19. Kinetics of the levels of anti-SARS-CoV-2 IgM and IgG antibodies over time. A serum sample was considered positive when the reaction intensity had a value equal to or above 30. Colored dots indicate individuals who became reinfected over the course of the study. **(A, B)** Show the reactivity of IgM and IgG specific for SARS-COv-2 antigens over time. The frequency of seroconversion is shown at the top of the figure. The number of individuals tested (*n*) varies according to the time point evaluated and is indicated on the graph. *p<0.05; **p<0.01; ***p<0.001 ****p <0.0001. Statistical significance was measured using a Kruskal-Wallis test at a significance level of 5%. **(C, D)** Each line represents one participant (*n* = 52). Dashed lines represent participants infected once with COVID-19. Reinfected participants are represented by solid lines. Red lines indicate the period before and after the reinfection. **(A-D)** Values below 30 are shown in the gray zone of the graphs. **(E, F)** Mean IgM and IgG immunoglobulin levels over time. For the analysis, the peak antibody level for each participant, observed from the seventh to the thirtieth day, was considered (30). Follow-up evaluation of anti-SARS-CoV-2 IgM and IgG levels was performed at days 60, 90, 180, 270, and 450 after enrollment in the study. A test was considered positive when the detected value was equal to or greater than 30.

**Figure 4:**
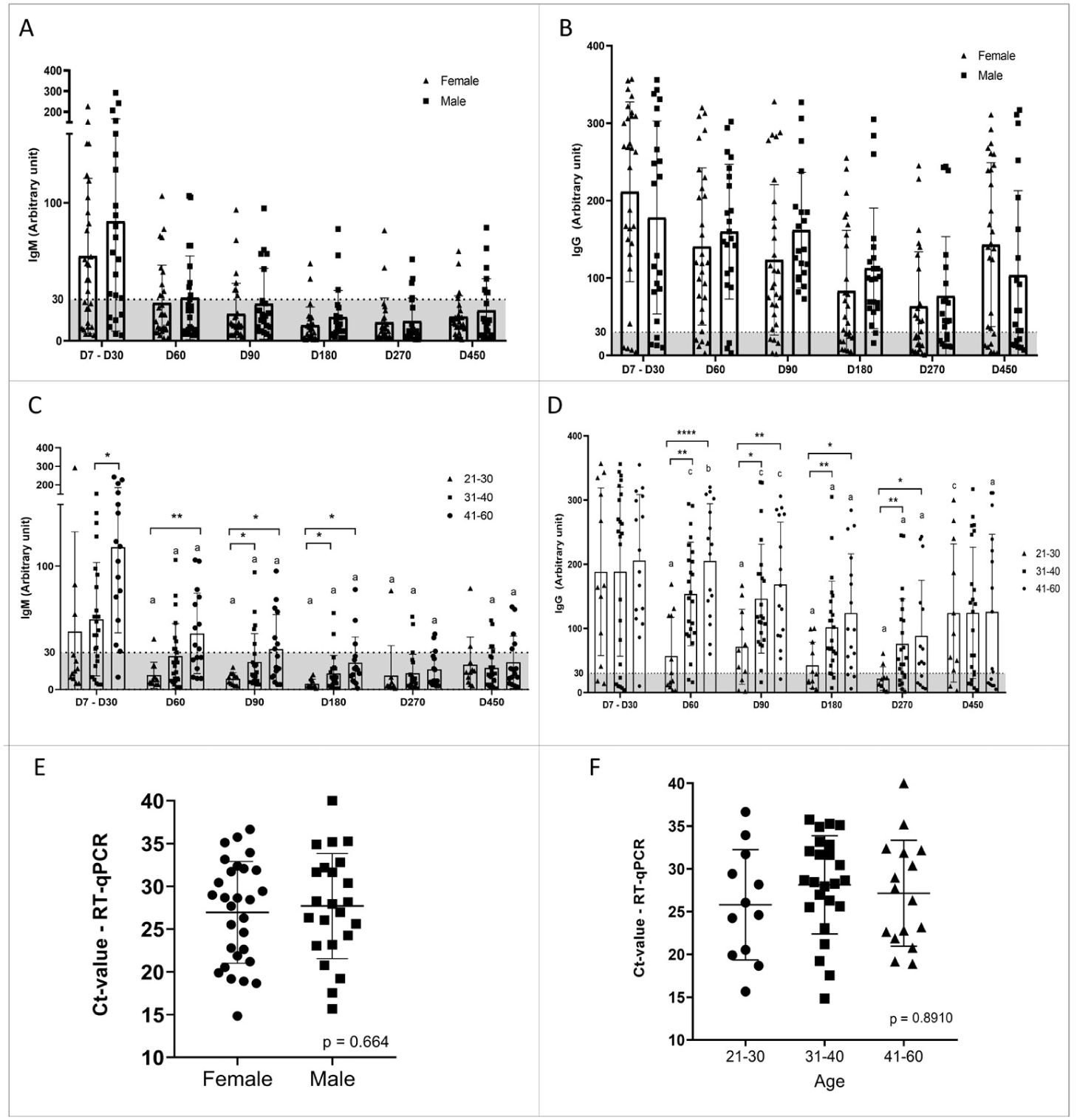
Dynamics of IgM and IgG levels against SARS-CoV-2 and Ct values over time stratified by sex and age. (**A-D)** The left and right panels represent the distribution of IgM and IgG levels, respectively. A serum sample was considered positive when the reaction intensity had a value equal to or above 30. Values below 30 are shown in the gray zone of the graphs. *P* values were determined using a Sidak’s multiple comparisons test, ANOVA, or unpaired *t*-test. **(A, B)** Stratification of the humoral response over time by sex. The triangles represent the antibody level of women and the squares that of men. There was no statistical difference. **(C, D)** Stratification of humoral response over time by age, 21-30 years (triangles), 31-40 years (squares), and 41-60 years (dots). “a” significant difference in comparison with D7-D30 (p<0.005); “b” significant differences in comparison with D180 (p< 0.005); “c” significant differences in comparison with D270 (p< 0.005); *p<0.05; **p<0.01; ***p<0.001 ****p <0.0001. **(E)** Ct values of males (squares) and females (dots). Statistical significance was measured using an unpaired *t*-test at a significance level of 5%. **(F)** Ct values in participants stratified by age: 21-30 years (dots), 31-40 years (squares), and 41-60 years (triangles). Statistical significance ANOVA at a significance level of 5%.

The dynamics of IgM and IgG levels over time are shown in Figures 3E and 3F. For IgM and IgG immunoglobulins, there were two behaviors: the drop in the mean antibody levels up to 180 and 270 days, respectively, followed by a slight increase after this time point. Tables 2 and 3 demonstrate the behavior of IgM and IgG estimated by the segmented model. The GEE results indicate that the average IgM and IgG values decreased by 0.606 and 0.645 units each day until days 180 and 270, respectively. After this time, the slope is positive for both IgM (0.363) and IgG (0.224). As shown in Figure 3E, the observed mean IgM levels were below the cutoff despite this upward trend. We can also observe that 60 days after symptom onset, the mean anti-SARS-CoV-2 IgM levels are below the cutoff, while anti-SARS-CoV-2 IgG levels drop, but not below the cutoff, around 270 days after symptom onset (Figures 3E and 3F).

**Table 2:**
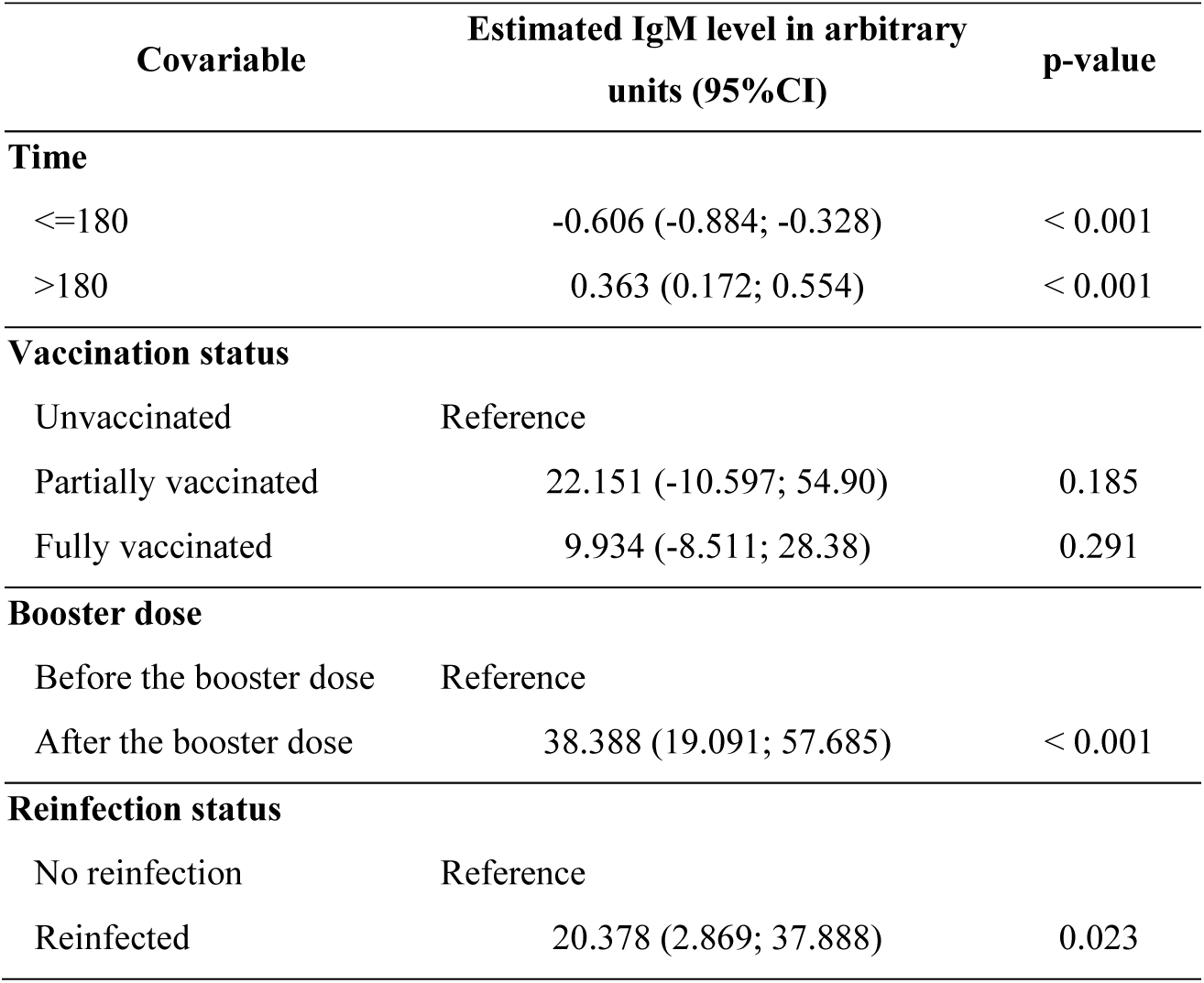
Parameter estimates according to Generalized Estimating Equations (GEE) analysis of the relative numeric scale of IgM levels. Vaccine status, follow-up time, booster dose, and reinfection were used as covariates.

| Covariable | Estimated IgM level in arbitrary units (95%CI) | p-value |
| --- | --- | --- |
| **Time** |  |  |
| <=180 | -0.606 (-0.884; -0.328) | < 0.001 |
| >180 | 0.363 (0.172; 0.554) | < 0.001 |
| **Vaccination status** |  |  |
| Unvaccinated | Reference |  |
| Partially vaccinated | 22.151 (-10.597; 54.90) | 0.185 |
| Fully vaccinated | 9.934 (-8.511; 28.38) | 0.291 |
| **Booster dose** |  |  |
| Before the booster dose | Reference |  |
| After the booster dose | 38.388 (19.091; 57.685) | < 0.001 |
| **Reinfection status** |  |  |
| No reinfection | Reference |  |
| Reinfected | 20.378 (2.869; 37.888) | 0.023 |

**Table 3:** Parameter estimates according to Generalized Estimating Equations (GEE) analysis of the relative numeric scale of IgG levels. Vaccine status, follow-up time, booster dose, and reinfection were used as covariates.

| Covariable | Estimated IgG level in arbitrary units (95%CI) | p-value |
| --- | --- | --- |
| **Time** |  |  |
| <=270 | -0.645 (-0.811; -0.479) | < 0.001 |
| >270 | 0.224 (0.127; 0.321) | < 0.001 |
| **Vaccination status** |  |  |
| Unvaccinated | Reference |  |
| Partially vaccinated | 15.977 (-15.343; 47.298) | 0.317 |
| Fully vaccinated | 26.848 (-7.077; 60.773) | 0.121 |
| **Booster dose** |  |  |
| Before the booster dose | Reference |  |
| After the booster dose | 52.221 (15.406; 89.035) | 0.005 |
| **Reinfection status** |  |  |
| No reinfection | Reference |  |
| Reinfected | 9.715 (-10.566; 29.996) | 0.348 |
| **Interaction** |  |  |
| Booster dose - Reinfection | 87.8 (27.3; 148.0) | 0.001 |
| Reinfection - Booster dose | 130.3 (82.19;178.30) | 0.001 |

No significant difference in IgM and IgG antibody levels with regard to the vaccination status of the participants was observed (Tables 2 and 3). However, after the booster dose, the anti-SARS-CoV-2 IgM and IgG levels both significantly increased. The IgM levels after the booster dose were 38.388 units higher. In addition, IgM levels were significantly higher after reinfection by 20.378 units. The interaction between these two covariables was not significant. In contrast, when we analyzed the IgG levels, the interaction between the booster dose and reinfection was significant, with the mean IgG levels 130.3 [95%CI: 82.19-178.30] units greater in those that received the booster dose and got reinfected. Before the booster dose, there was no difference in the mean IgG level when comparing reinfected and non-reinfected participants (p = 0.348). After the booster dose, the mean level of IgG for reinfected individuals was 87.8 [27.3; 148.0] units, which was higher than for non-reinfected individuals. For non-reinfected individuals, after the booster dose, the mean IgG was 52.22 units higher than before.

Of the 19 participants who got reinfected (Figure 5), 13 (68.4%) were after and 5 (31.6%) before the booster dose. In four (21.0%), increased levels of IgM and IgG antibodies were not observed, even after booster dose and re-infection, these individuals being non-reactive to SAR-CoV-2 antigens at day 450 for both immunoglobulin classes (Figure 5: F, G, J, O). One individual (Figure 5: F) did not seroconverted during the whole period of follow-up. Of the remaining 15 individuals who got reinfected, considering IgM dynamics, four of them never seroconverted during the follow-up period (Figure 5: B, H, K, S). Four individuals (Figure 5:C, D, E, R) who were negative at baseline only had IgM seroconversion after reinfection or the booster dose. For four other individuals (Figure 5: L, M, P, Q) that were IgM reactive at baseline, but became non-reactive during the longitudinal assessment, the booster dose and reinfection were not able to stimulate production of this class of antibody. For the remaining three participants (Figure 5: A, I, N), IgM levels declined to non-reactive, but the booster dose or reinfection seroconverted them again. For IgG, the dynamics are different, since all 15 reinfected participants seroconverted at some point during the study. Of these, one only seroconverted after the booster dose (Figure 5: K). The other 14 participants had reactive IgG antibodies at baseline. For all of them, anti-SARS-CoV-2 IgG antibody levels declined, with seven participants becoming non-reactive to SARS-CoV-2 antigens for this class of antibody. For the remaining seven seroreactive participants, six showed increased IgG levels after reinfection or booster dose (Figure 5:H, I, L, M, P, R). For the individuals who became IgG seronegative over the course of the study, the booster dose or reinfection was able to cause secondary IgG seroconversion of all such participants (Figure 5: A, B, C, D, E, N, S).

**Figure 5:**
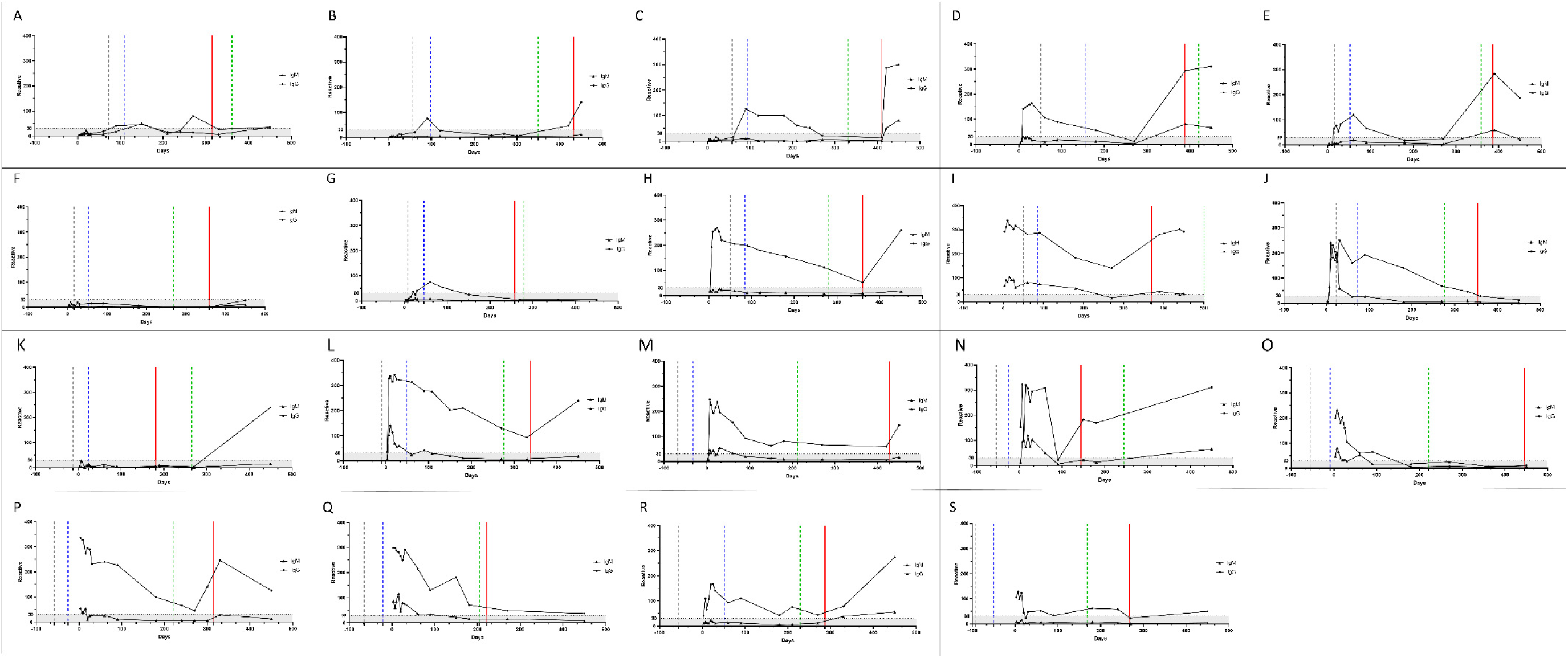
IgM and IgG antibody profiles of participants who were reinfected. **(A-S)** The triangles represent the IgM and the dots the IgG values. The gray, blue and green dashed lines represent the beginning date of the vaccination, the complete vaccination scheme, and the booster dose, respectively. The red line represents the reinfection period. A serum sample was considered reactive when the reaction intensity had a value equal to or above 30. Values below 30 are shown in the gray zone of graphs.

### 2.3. Reinfection

Every 30 days or when participants presented symptoms that indicated a suspicion of COVID-19 reinfection, saliva or naso-oropharyngeal swab collection was performed to detect viral RNA. At the same time, a capillary blood sample was collected to evaluate immunoglobulins. Of the 52 participants, 46 (88%) described at least some symptoms compatible with suspected COVID-19, with 9 (17%) reporting it once, 19 (36%) twice, 9 (17%) three times, five (10%) four times, and four (8%) participants reporting five times. In total, although 114 suspected episodes of COVID-19 were recorded, only 19 were confirmed by RT-qPCR. Of all the symptom episodes/types (*n* = 185) reported by participants during the study, cough (61.6%) followed by congestion or runny nose (49.2%) and sore throat (41.1%) were the most frequent. The most common symptoms reported during the suspected and confirmed reinfections were congestion or runny nose (57.0% and 74%), cough (40.4% and 63%), and sore throat (45.6% and 47%), respectively. The profile of symptoms presented by reinfected individuals differs from those reported during the first SARS-CoV-2 infection, with statistical significance (p<0.001) for headache, fever and chills, myalgia, anosmia, and ageusia between groups were observed (Table 4). Ageusia and anosmia were not reported in confirmed cases of reinfection (p<0.001). Other symptoms reported during the first infection in the suspected and confirmed cases of reinfection can be seen in Figure 6A and Table 4.

**Figure 6:**
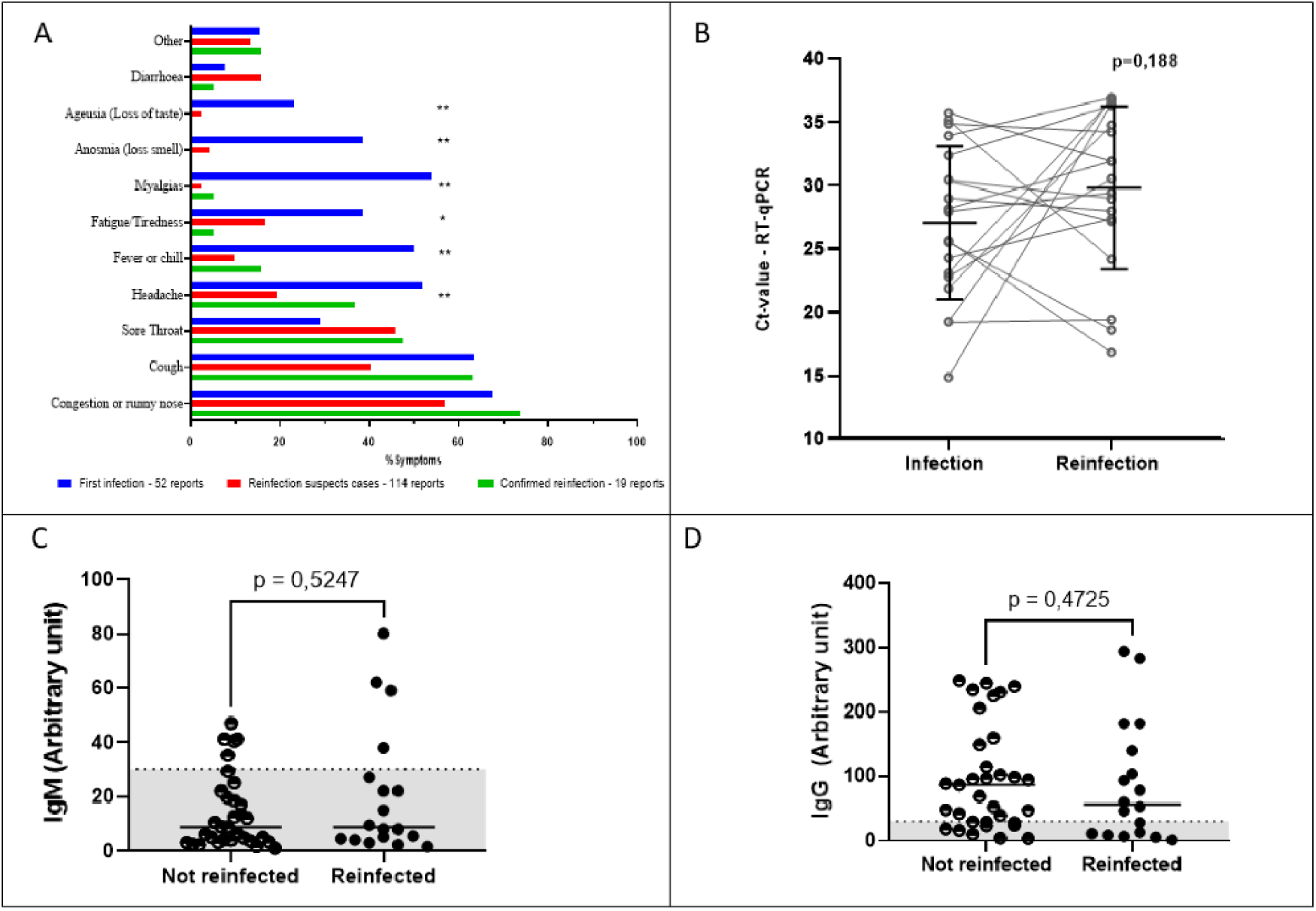
Main self-reported symptoms, dynamics of IgM and IgG levels, and Ct-values stratified according to SARS-CoV-2 infection/reinfection status. **(A)** Main self-reported symptoms described by participants. Blue bars indicate the percentage of symptoms reported by the 52 COVID-19 positive participants. Red bars show the symptoms reported in cases of suspected reinfection, but without confirmation by detection of viral RNA. Green bars show the main symptoms of participants with confirmed reinfection by SARS-CoV-2. Statistical significance was measured using either the Chi-square test or Fisher’s exact test **p < 0.001, *p=0.002. **(B)** SARS-CoV-2 viral load. Statistical significance was measured using a paired *t*-test at a significance level of 5%. **(C, D)** Antibody levels of reinfected and non-reinfected participants. For individuals who were not reinfected, the values shown are the arithmetic means of multiple measurements of their circulating antibody levels taken throughout only the period of the study when the waves of the Delta and Omicron variants occurred. For individuals who were reinfected, the antibody levels shown are single point estimates determined only at the time of reinfection confirmation. The left and right panels represent the distribution of IgM **(C)** and IgG (**D)** levels, respectively. A serum sample was considered positive when the reaction intensity had a value equal to or above 30. Values below 30 are shown in the gray zone of graphs. P values were determined using the Mann-Whitney *U* test at a significance level of 5%.

**Table 4.** Main symptoms reported by the final study cohort.

| Symptoms | First infection | Suspected reinfection | Confirmed reinfection | Total | p-value |
| --- | --- | --- | --- | --- | --- |
| Congestion or runny nose | 35 (67.3%) | 65 (57.0%) | 14 (73.7%) | 114 (61.6%) | 0.234 |
| Cough | 33 (63.5%) | 46 (40.4%) | 12 (63.2%) | 91 (49.2%) | 0.010 |
| Sore throat | 15 (28.8%) | 52 (45.6%) | 9 (47.4%) | 76 (41.1%) | 0.106 |
| Headache | 27 (51.9%) | 22 (19.3%) | 7 (36.8%) | 56 (30.3%) | <0.001 |
| Fever or chill | 26 (50.0%) | 11 (9.6%) | 3 (15.8%) | 40 (21.6%) | <0.001 |
| Fatigue/Tiredness | 20 (38.5%) | 19 (16.7%) | 1 (5.3%) | 40 (21.6%) | 0.002 |
| Myalgias | 28 (53.8%) | 3 (2.6%) | 1 (5.3%) | 32 (17.3%) | <0.001 |
| Anosmia (loss smell) | 20 (38.5%) | 5 (4.4%) | 0 (0.0%) | 25 (13.5%) | <0.001 |
| Ageusia (loss of taste) | 12 (23.1%) | 3 (2.6%) | 0 (0.0%) | 15 (8.1%) | <0.001 |
| Diarrhea | 4 (7.7%) | 18 (15.8%) | 1 (5.3%) | 23 (12.4%) | 0.287 |
| Other<sup>1</sup> | 8 (15.4%) | 15 (13.2%) | 3 (15.8%) | 26 (14.1%) | 0.855 |
| Total | 52 (28.1%) | 114 (61.6%) | 19 (10.31%) | 185 (100%) |  |

1- Other symptoms reported to healthcare professionals included: malaise, throat irritation, pain in the face, abdominal pain, inappetence, asthenia, arthralgia, retro orbital pain and sweating, nausea or vomiting, and red or irritated eyes.

The first five (26%) cases of reinfection occurred between August and September 2021 (Table 5). At this time, the Delta and Gamma variants were circulating in Belo Horizonte (Figure 7). Between December 2021 and January 2022, 12 cases (63%) of reinfection were confirmed, a period that overlaps with the new wave of transmission caused by the Omicron variant (Figure 7). Of the reinfection samples collected during this period that were sequenced, nine were identified as the Omicron variant. Sequencing samples with low viral load (Ct value > 36) was not possible. The mean CT of infected participants was 30.17 (± 6.5) (Table 5). There was no significant difference between Ct values detected during infection and reinfection (Figure 6B). When assessing the humoral response of reinfected and non-reinfected individuals, we did not observe differences in IgM and IgG levels between these groups (Figures 6C and 5D). The intervals between infection and vaccination and seroconversion status are given in Tables 5 and 6.

**Figure 7:**
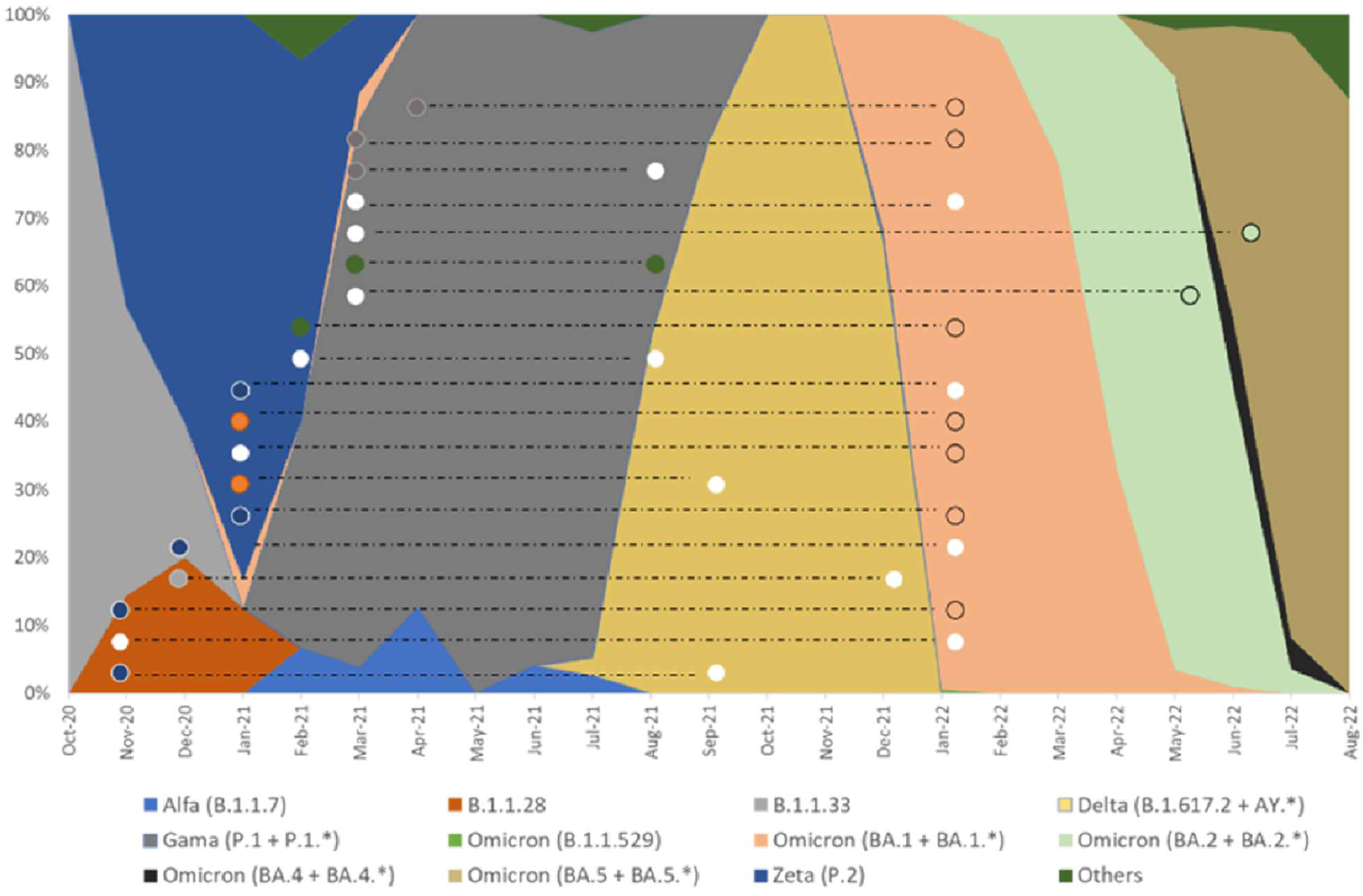
Overlap of genomic strains sequenced in this study in relation to strains circulating in Belo Horizonte during the same period. Absolute frequency of SARS-CoV-2 genomic samples sequenced in Belo Horizonte from October 2020 to August 2022 (Stacked area graph). The colors inside the circles indicate the sequence of SARS-CoV-2 strains. White dots represent samples for which the SARS-CoV-2 strain was not determined. The dashed lines connect results obtained in samples from the same participant during their first infection and subsequent reinfection

**Table 5:**
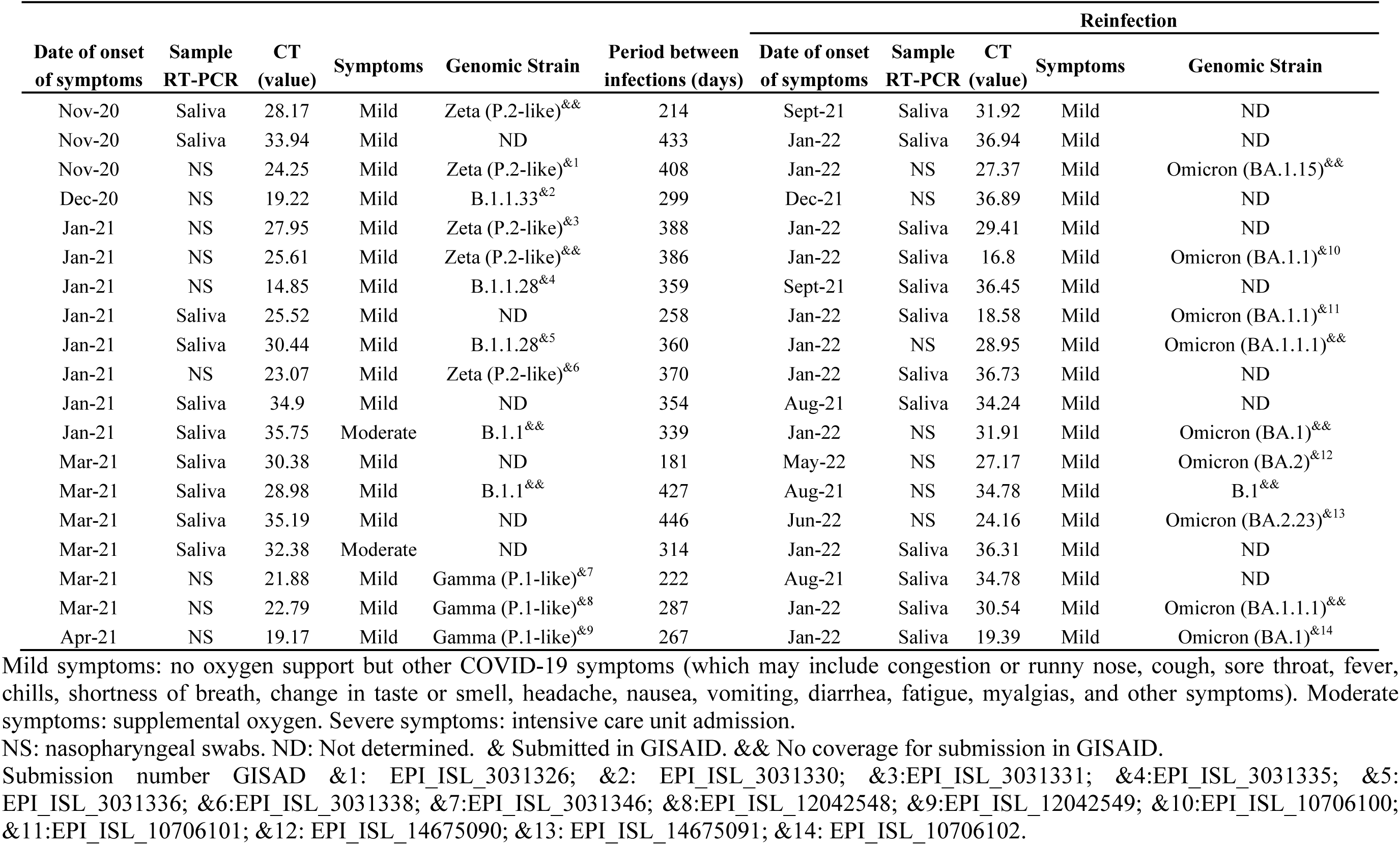
Genomic strain, Ct value, and classification of symptoms of confirmed cases of SARS-CoV-2 infection and reinfection.

**Table 6:**
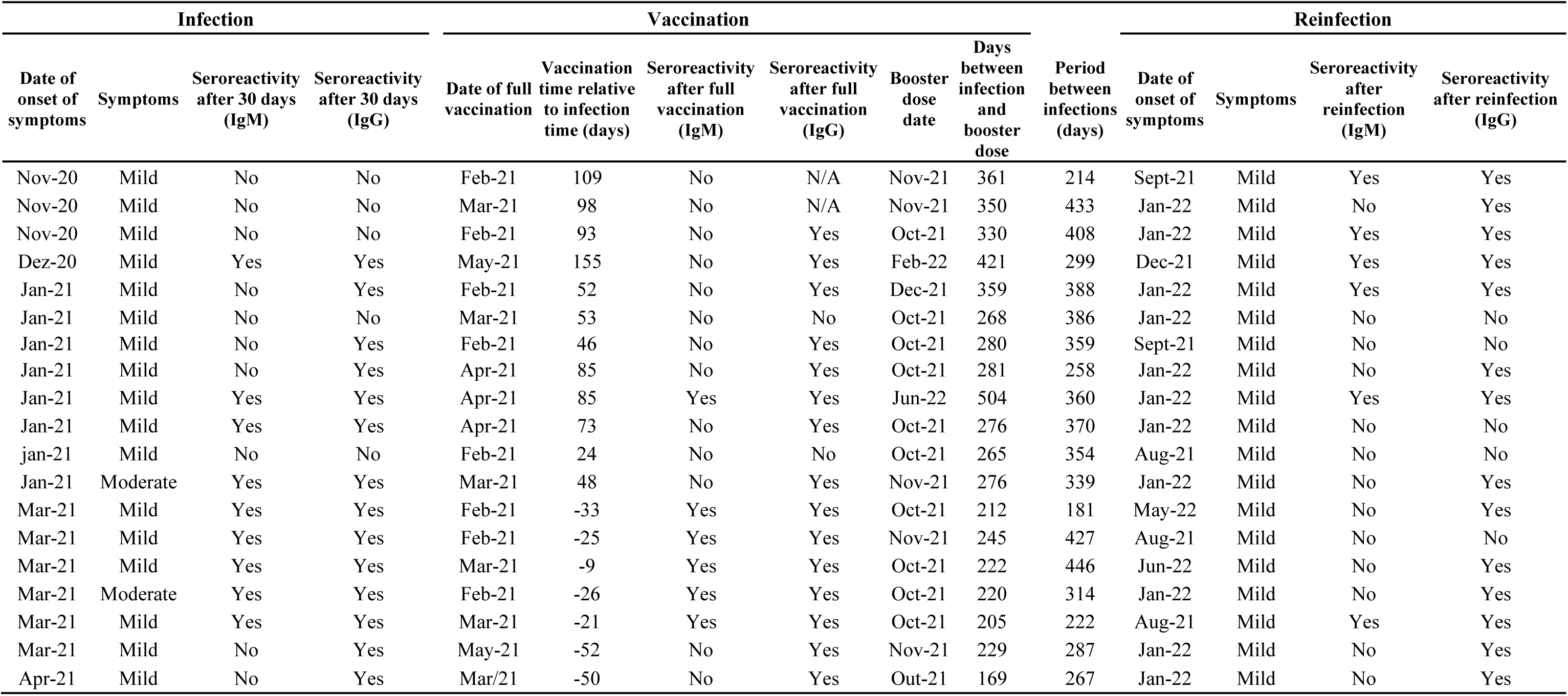
Seroconversion of IgM and IgG after infection, vaccine doses, and reinfection.

## 3. Discussion

Elucidating the kinetics of the humoral response to SARS-CoV-2 is crucial for controlling the pandemic, and for designing, planning and implementing the most appropriate vaccine schemes (1). Here, we investigated for 450 days the dynamics and longevity of IgM and IgG antibodies from healthcare professionals. The long follow-up period allowed us to monitor dynamics of the humoral response after vaccination booster doses and reinfection.

Many previous studies have evaluated this profile. However, such studies have a variety of follow-up durations ranging from 50 days (26), 100 days (27–30), 210 days (31), 360 days (32), and 480 days (33). Some of these studies followed specific antibodies against the nucleocapsid (N), the receptor-binding domain (RBD), or the spike (S) protein, using either ELISA assays, immunochromatographic tests, or looking for neutralizing antibodies. In our study, a commercial immunochromatographic test was used to detect both IgM and IgG against viral proteins. It has been shown by some authors that there is a correlation between the detection of total and neutralizing antibodies (29, 33, 34).

Our study reinforces the view, and provides evidence, that antibodies are initially produced, but decline over time. The drop of IgM starts on the 30th day, reaching non-reactive levels by the 60th day after symptom onset, the same profile as previously described (29, 30). For IgG, a less pronounced decline is observed, and around the ninth month, we observe the lowest mean reactivity. The stability of the IgG reactivity for three months has been previously demonstrated (29, 30). Gil-Manso et al. (2020)(35) and Gaebler (2021) (36) observed that the IgG response lasted longer, about six months, and the levels of neutralizing activity were proportional to anti-RBD IgG antibody titers. Some studies demonstrate that the duration of the response depends on the studied target: anti-RBD antibodies remained stable for between six to 12 months, while anti-N antibodies decreased over the same period (32). In contrast, Yang (2022)(33) described a peak of anti-RBD antibodies around 120 days after the onset of symptoms with a subsequent decline, maintaining positivity until day 400 after symptom onset. A recent study describes that for non-vaccinated individuals, IgG antibodies, evaluated by ELISA, persist for one year (37).

The results of longitudinal studies, however, may critically vary according to different conditions and variables. Mioch et al. (2023)(37) report that loosening epidemiological control measures increases the chance of re-exposure to the virus. The sensitivity of the tests and different methods used may vary. In addition, patients with comorbidities may have different antibody kinetics, as demonstrated by the rapid decline of antibodies in diabetic patients (38). Yang et al. (2022)(33) conducted a long follow-up of the humoral response in individuals confirmed to be free of re-exposure and vaccination against SARS-CoV-2. Although their study is significant, it does not portray the global reality since we have more than 60% of the population vaccinated worldwide, while in Brazil more than 80% of people completed the vaccination schedule and almost 50% took a booster dose (1). In our cohort, all healthcare professionals who worked on the front line completed the vaccination schedule with the Coronavac vaccine and took the Pfizer vaccine as a booster dose (6)

For our analyses, we considered the peak of reactivity to be between days 7-30 (D7-D30). Thus, there was no difference in the increase of antibody levels for fully and partially vaccinated and unvaccinated individuals. Based on the dynamics presented, the slight increase observed in the mean antibody levels (D60-D90) may have been induced by the Coronavac vaccination (39). After using the Generalized Estimating Equations (GEE), it is possible to establish that the Pfizer booster dose increased antibody levels for both IgM and IgG. The decline of antibodies after the Coronavac vaccination, even in individuals who became infected, has also been already described. The booster dose is essential to restimulate the humoral response (40–42) and the same profile is also observed for other vaccines (43–46).

Vaccines are essential to reduce the morbidity and mortality from COVID-19. However, they cannot completely prevent new infections and reinfections (43). In our cohort, 36% of participants were reinfected either before (31.6%) or after (68.4%) the booster dose. The reinfections coincided with the spread of new variants in Brazil, such as Delta and Gama in August-September 2021, and Omicron in December 2021. The emergence of new variants has been a matter of great concern, as they can reduce neutralization and even escape vaccination, as demonstrated for Delta (39, 47), Beta and Gamma (40, 48), and Omicron (49–51) variants.

Cases of reinfection have been reported since the beginning of the pandemic (12, 20, 52–54), including in Brazil (55–57). Studies have shown that an acquired immune response can reduce the risk of transmission by up to 90%, with an interval of 6-10 months (58–60). The reinfection rate is relatively low, ranging from 0.1-0.65(18–20, 61–63), with the highest rates reported in the UK study at 1.9% and 4.5% in India (24). In our study, we reported a high rate of reinfection that might be due to some important factors. Most of the reinfections occurred nine months after the first infection, when antibody levels were already low. Additionally, reinfections coincided with the entry of new variants into Brazil, which, as already mentioned, have a high rate of transmission, and escape from immune responses (14, 21, 25, 60, 64).

Some studies show that men are more likely to test positive and develop severe COVID-19 (65). Petersen et al., (2022)(66) showed a faster decrease in IgG in males. Frauke et al. (2022) (34)observed that the decline of neutralizing antibodies in men was faster than in women but that afterward, there was no difference in response. The same was observed by (67), in which no correlation exists between neutralizing antibodies and biological sex. We also did not notice any difference in the behavior of the humoral response between men and women. Evaluating reinfection cases, our data corroborate the data of Alexander Lawandi et al., (2022)(68), in which we observed a higher rate of reinfection in women, which contrasts with the review made by Sahar (2022)(24), while other studies did not find a relationship between sex and reinfection(69). This heterogeneity of results demonstrates that other factors must be evaluated, such as comorbidity, lifestyle, workplace, biological and immune differences. Understanding sex differences is fundamental to improving disease management, predicting outcomes, and planning specific interventions for men and women.

There is no relationship between age and cases of reinfection in health workers in our study, as demonstrated by Alejandra Svartza (2023)(70). There was also no difference between the general population and healthcare workers who were reinfected. Ren et al., (2022)(22) and Sahar Ghorbani (2022)(19) review that there is a wide age distribution among reinfected patients, ranging from 15 to 99 years. Hansen (2021)(71) described that an age greater than 65 years might influence the increase in the relative risk of reinfection. Healthcare professionals in our cohort who became reinfected were between 24 and 55 years old, and all of them had mild symptoms. Even though age was not associated with reinfection, we observed that the dynamics of antibody levels varied over time and behaved differently across age groups. The level of antibodies produced by the younger group was lower than the older groups, as observed by others works (66, 72, 73)

Many studies have sought to understand and relate antibody profiles and Ct values with disease severity and protection against reinfection (27, 33, 74, 75). Omid Dadras et al., 2022 (76), concluded that the relationship between viral load and disease severity is inconclusive. We observed no significant difference in viral load at first infection and reinfection. Except for two participants who required oxygen support for their first infection, all other participants had mild disease. As expected, the main symptoms reported were nasal congestion, coughing, sore throat, and headache. Other symptoms were also reported as described in other studies (22). The reinfection period of most participants in our study overlaps with the spread of the Omicron variant, as described by Menni et al (2022), Karina Vihta (2022)(77, 78) and Machado Curbelo (2022)(79), who demonstrated that anosmia and ageusia were less frequently associated with the Omicron variant. No reinfected participants reported anosmia or ageusia in our study. Symptoms such as cough, fever, shortness of breath, myalgia, fatigue, and headache were less frequently reported by participants who had suspicion of COVID-19 either confirmed or not by RT-qPCR. In contrast, sore throat was more frequently reported by those participants, although no significant difference in frequency was observed among infection/reinfection status. These clinical conditions corroborate those described by Karina Vithta (2022)(78).

Our study has some limitations. We evaluated total antibodies and did not determine if they were neutralizing, nor the quality of memory B cells necessary to produce antibodies against reinfection. Also, we monitored reinfection in symptomatic participants, but there is a possibility that cases of asymptomatic reinfection also occurred (24).

The humoral response declines after the first infection and vaccination but increases substantially after reinfection and booster doses, especially for the IgG antibody class. There is no association between circulating antibody levels and cases of reinfection. Overall, we demonstrated that even after the booster dose, health professionals can become reinfected with new variants of SARS-CoV-2. Therefore, our study demonstrates that prolonged protection after COVID-19 infection, even after the booster dose, does not prevent reinfection by new variants, which contrasts with the prolonged immune response cited by other studies (66, 80). Studies show that previous infection and booster dose reduce the risk of reinfection, as seen in Switzerland (81), Qatar (61), and United States of America (82). However, studies have shown a significant decrease in the effectiveness of the vaccine against the Omicron variant within a few months after administration (83). Although our study was not designed to assess whether the booster dose would prevent reinfection, the evidence suggests that the booster dose was not effective in preventing reinfection. Those differences might have been impacted by the vaccine scheme and/or type used during the vaccination campaign.

In this context, our data reinforce the importance of robust surveillance in viral genomics and in the immune response of individuals, especially in high-risk individuals, such as the immunocompromised and health care professionals. Investing in these tools is essential for preparing and responding to new variants and future pandemics.

## 4. Materials and Methods

### 4.1. Ethical approval

This study was conducted in accordance with current legislation including the Declaration of Helsinki and Resolution No. 466/2012 of the Conselho Nacional de Saúde do Brasil. Ethical approval was obtained from the institutional review board of the Instituto René Rachou, Fundação Oswaldo Cruz, CAAE: 31.919520.8.0000.5091, approval numbers: 4177931; 4291836; 4343318; 4624187; 5294423. Written informed consent was obtained from all participants before any study procedure was undertaken.

### 4.2. Study population and enrollment

The population selected for this study was composed of HCW who worked in at least one of the public hospitals: Hospital das Clínicas (HC) at the Universidade Federal de Minas Gerais, the Unidade de Pronto Atendimento (UPA) Centro-Sul, and the Hospital Metropolitano Dr. Célio de Castro (HMDCC). All three health centers are in the city of Belo Horizonte, the state of Minas Gerais, Brazil. As inclusion criteria, in addition to what has already been mentioned above, all participants had to (i) present with at least one of the following symptoms within the previous seven days: fever (equal to or greater than 37.5°C), cough (dry or productive), fatigue, dyspnea, sore throat, anosmia/hyposmia, and/or ageusia; and (ii) have a positive result for the detection of SARS-CoV-2 RNA by RT-qPCR. Participants were invited by telephone, and those who met the inclusion criteria were included in the study. Individuals who reported volunteering in COVID-19 vaccine clinical trials, prior diagnosis of COVID-19, or reported pregnancy, were excluded. Hospitalization also resulted in loss of follow-up due to inability to perform the tests. The enrollment of participants took place between October 2020 to April 2021. Individuals were followed up for 450 days, with capillary blood samples collected on days 7, 10, 15, 20, 25, 30, 60, 90, 180, 270, and 450 after being enrolled in the study (referred to as D7, D10, D15 and so on). In addition to capillary blood collections, participants were contacted every 30 days after the 60th day (from December 2020 to June 2022) to assess possible SARS-CoV-2 reinfection. The predetermined symptoms which were monitored as evidence for reinfection were: fever (equal to or greater than 37.5 °C), cough (dry or productive), sore throat, fatigue, dyspnea, and diarrhea. Also, any suspicious symptom established by medical criteria was monitored. In January 2021, during the study period, public roll-out of the vaccination scheme using the CoronaVac vaccine was started in Brazil. Fifteen days after the second dose, the initial two-dose vaccination scheme is considered “complete”. However, in September 2021, a booster dose was rolled-out (6). The workflow for our study is shown in Figure 1.

### 4.3. Study design

All symptomatic healthcare professionals were tested for SARS-CoV-2 by RT-qPCR using the Charite Institute protocol (84). During the isolation period, kits were sent to participants to perform a self-collection of saliva or nasopharyngeal and oropharyngeal swabs on D1, D3, and D5 after enrollment in the study. In addition to the material for saliva/swab sample collection, the kits included materials, together with an instruction manual, for performing a rapid test to detect IgM and IgG antibodies against SARS-CoV-2 antigens (TR DPP® COVID-19 IgM/IgG - Bio-Manguinhos). Participants characterized as infected with SARS-CoV-2 must have had at least one positive result for detection of SARS-CoV-2 RNA on D1, D3 and/or D5. Participants characterized as non-infected presented negative results in all three samples analyzed, and were not included in the follow-up period of the study.

### 4.4. RNA extraction

RNA was extracted from the nasopharyngeal and oropharyngeal swabs or saliva samples using the QIAamp® Viral RNA kit (QIAGEN®, USA) according to the manufacturer’s instructions. Briefly, 140 μL of sample was added to 560 μL of AVL buffer containing carrier RNA, and after 10 minutes at room temperature, 560 μL ethanol were added. This solution was applied to an RNA affinity column and this column was centrifuged at 6,000 x g for 1 minute. Then, the column was washed with AW1 and AW2 buffer solutions in that order. After the washing process, RNA was eluted using AVE solution and used in the RT-qPCR test.

### 4.5. RT-qPCR (quantitative PCR)

The RT-qPCR reactions were performed using the ViiA™ 7 Real-Time PCR System of the communal Real-Time PCR platform at the Instituto René Rachou. The RT-qPCR assays were performed using 5 uL of sample RNA, and the 200 nM GoTaq® Probe 1-Step RT-qPCR System Kit (Promega). This kit uses GoTaq Probe qPCR Master Mix with dUTP (10 uL), GoScript RT Mix for one-step RT-qPCR (0,4 uL), sense and antisense primers (400 nM) and nuclease-free water to a 20.0 uL final volume. The conditions for the amplification were: 45 °C for 15 minutes and 95 °C for 2 minutes, followed by 40 cycles of denaturation at 95 °C for 15 seconds and hybridization at 60 °C for 1 minute. The results were analyzed using the Thermo Cloud platform, according to the following criteria: samples with amplification of the *E* gene (Ct<37) and the RNAse *P* gene (RP) (Ct<35) were considered positive; samples without *E* gene amplification or with detection above Ct 37, with RP amplification (Ct<35), were considered negative. Samples with RP amplification above Ct 35 were considered invalid, and the test was performed again using RNA obtained from another extraction of the samples collected.

### 4.6. TR DPP® COVID-19 IgM/IgG test (Bio-Manguinhos)

In order to detect IgM and IgG antibodies against SARS-CoV-2, the DPP® COVID-19 IgM/IgG kit supplied by Bio-Manguinhos (FIOCRUZ, Brazil) was used according to the manufacturer’s instructions. Briefly, a digital puncture was performed, and a blood sample was diluted in the buffer provided in the kit. The sample was then applied to the cassette. After 5 minutes, 9 drops of the buffer were added to the cassette, and the results were read after an additional ten-minute period. The interpretation of the test was performed with the aid of the DPP® Micro Reader, which provides the intensity of the reactive line. Values equal to or greater than 30 for the IgM and IgG antibodies were considered positive. In order to assess antibody levels, the test was performed on scheduled days and/or when the participant was suspected of reinfection (Figure 1).

### 4.7. Next-generation sequencing

Positive samples, with cycle threshold (Ct) lower than 36, were sequenced by Next-Generation Sequencing (NGS) on the Illumina MiSeq Platform using the Illumina COVIDSeq Kit for library construction (Illumina, San Diego, USA) generating paired-end reads 150 bp long. The raw reads were trimmed using Trimmomatic version 0.39 (85) with a sliding window of 4 nucleotides with a minimum average Phred score of 20. Trimmed reads smaller than 50 bp were removed. The filtered reads were mapped to the SARS-CoV-2 reference genome (NC_045512) using BWA version 0.7.17 (Li 2009) with the default parameters. The nucleotide variants were identified using iVar version 1.3. (86), with a minimum frequency of 40% and depth of 30 reads. The consensus sequences generated by iVar were submitted to Pangolin version 4.2 (87) to identify the coronavirus lineage. Sequences that met the GISAID criteria were submitted to the EpiCoV database.

### 4.8. Statistical analysis

GraphPad Prism v.8.0.1., Jamovi 2.3.18.0 (https://www.jamovi.org/), and the statistical software R (https://www.r-project.org/) version 4.1.2 (88) were used for data analysis and generation of figures. Data organization and pre-processing of some graphs and figures were done using Excel and PowerPoint (Microsoft 365). The chi-square test or the Fisher’s exact test were used to assess the association of categorical demographic variables and the infection status of individuals, as well as the correlation of symptoms reported during the first infection, reinfection and in suspected cases of reinfection. Paired and unpaired *t*-tests were used to analyze the difference in viral load between infection vs reinfection, and females vs males, respectively.

Generalized Estimating Equations (GEE) (version 1.3.9) were used to evaluate the longitudinal data. Proposed by Lian and Zeger (1986)(89), the proposed model jointly estimates an average effect and intra-individual variations, considering the structure of correlation or dependence between the repeated measures. The outcome of interest was the numeric scale of IgM and IgG levels and the covariates were vaccine status, follow-up time, booster dose and reinfection. In the GEE model, the peak of IgM and IgG levels in the interval from D7 to D30 was considered. A model segmented in time, 180 and 270 for IgM and IgG, respectively, was adjusted to make the average structure more accurate.

In order to assess whether there is an association between antibody levels and reinfection, differences in the levels of IgM and IgG antibodies against SARS-CoV-2 among individuals with a confirmed case of reinfection and those not reinfected were evaluated. This analysis only considered samples taken during the period of the study when reinfection occurred (i.e., between July 2021 and July 2022), which was also the period when the delta and omicron variants circulated in Belo Horizonte. For this evaluation, the Mann-Whitney *U*-test was used. For individuals who did not become reinfected, the arithmetic means of all of the values of the antibody levels determined during the period stated above were considered. For individuals who became reinfected, we considered only the values of the antibody levels determined when reinfection was confirmed.

## Acknowledgments

The authors thank all participants for their involvement in the study, and the healthcare and administrative personnel at the hospitals participating in this study. We acknowledge the Program for Technological Development in Tools for Health–PDTIS–FIOCRUZ for the use of its Real– Time PCR and Next Generation Sequencing facilities; the Clinical Research Platform of Fiocruz for monitoring the study documents and for generating and managing the study database; the project support service of the IRR for project management. We also thank Edward José Oliveira, Marina Moraes Mourão and Roberta Lima Caldeira for their contributions towards the conceptualization of the study and Luke Baton for revising the language of the manuscript.

